# Fundamental parameters governing the transmission of conjugative plasmids

**DOI:** 10.1101/2023.07.11.548640

**Authors:** Jorge Rodriguez-Grande, Yelina Ortiz, M. Pilar Garcillan-Barcia, Fernando de la Cruz, Raul Fernandez-Lopez

## Abstract

Because their ability to spread infectiously, conjugative plasmids represent a major route for the propagation of antibiotic resistances. In the wild, there are hundreds of different plasmids, but their distribution, prevalence and host range are extremely variable. To understand the reasons behind these ecological differences, we need to obtain precise models for plasmid transmission dynamics. Here, we show that the dynamics of plasmid transmission follow a simple ecological model known as Holling’s Type II Functional Response. Plasmid transmission dynamics can be faithfully captured using two parameters: the searching rate (the pace at which a donor bacterium encounters a suitable recipient) and the handling time (the time required for a donor to transfer the plasmid). By analyzing the dynamics of different plasmid prototypes, we show that these parameters are characteristic of the plasmid transfer machinery. Quantifying the dynamics of plasmid spread may shed light on the epidemiology of antibiotic resistance genes.

## Introduction

In bacteria, a sustained encroachment of antimicrobial resistances is threatening our ability to treat infection. Antibiotic therapy is a safe and effective tool to fight bacterial pathogens, but its efficacy declines as the prevalence of antibiotic resistance genes (ARGs) increases. Bacterial conjugation is a key driving force in the spread of ARGs, which are frequently encoded in plasmids (Castañeda-Barba et al., 2024; Orlek et al., 2023). Conjugative plasmids transmit themselves from one cell to the other, in a process that resembles the spread of pathogens among susceptible populations (Anderson & May, 1979; Bergstrom et al., 2000; Hernández-Beltrán et al., 2021; Levin et al., 1979; May & Anderson, 1979; Simonsen et al., 1990). Obtaining quantitative models for plasmid infection dynamics is key, not only for microbiologists studying the mechanisms of conjugation, but also for epidemiologists analyzing the spread of antibiotic resistances.

Since plasmid transmission requires cell-to-cell contact between a donor (D) and a recipient cell (R), conjugation has been classically assumed to follow mass-action kinetics (Levin et al., 1979; Smets & Lardon, 2009; Zhong et al., 2012; Zwanzig et al., 2019). Seminal models posed that the rate of transconjugant (T) formation may be approximated as the product of a particular conjugation rate, γ, multiplied by the concentrations of [D] and [R] (Stewart & Levin, 1977). Such approximation is formally and conceptually identical to density-dependent models (DDT) for the transmission of infectious agents (Figure 1A). But the dynamics of plasmid conjugation are more complex than that of classical pathogens. In bacteria, the timescales of plasmid transmission and vegetative growth are of similar magnitude, thus new T cells may arise either by conjugation or by host replication (Bethke et al., 2023; Turner et al., 1998). Early models for plasmid conjugation, such as Simonsen’s End-Point Method, devised clever methods to deconvolve both contributions (Simonsen et al., 1990). More modern approaches included further complexities of plasmid propagation, such as the effect of population spatial structure, or the stochastic nature of plasmid conjugation (Fox et al., 2008; Huisman et al., 2022; Kosterlitz et al., 2022; Krone et al., 2007). Although the quantitative framework to study conjugation gained in complexity along the years, most models kept as axiomatic that the conjugation rate shall be proportional to D and R concentrations (Wan et al., 2011; Werisch et al., 2017; Zhong et al., 2010, 2012). While this can be logically drawn from first principles (conjugation cannot happen without physical contact), its validity along the natural range of bacterial densities cannot be taken from granted. It may be possible that the rate of encounters ceases to be a limiting factor as cells reach a certain concentration. This density limitation occurs in the transmission of some infectious agents (McCallum et al., 2009; O’Keefe, 2005). Many sexually transmitted infections do not follow DDT dynamics, but instead can be better modeled using frequency-dependent transmission (FDT, Figure 1B). In ecology, a similar situation occurs in predator-prey dynamics. Predation rates are limited, not only by the availability of prey, but also by the time required for the predator to consume its catch (Holling, 1959). This double dependence generates a biphasic regime, in which DDT becomes dominant at low prey densities, while FDT dominates at higher ones (Figure 1C). These dynamics can be modeled using Holling’s type II functional response, which smoothly captures this biphasic regime using two parameters: the searching rate (K_on_, which governs the DDT regime), and the handling time (τ, which sets the FDT limit).

**Figure 1.**
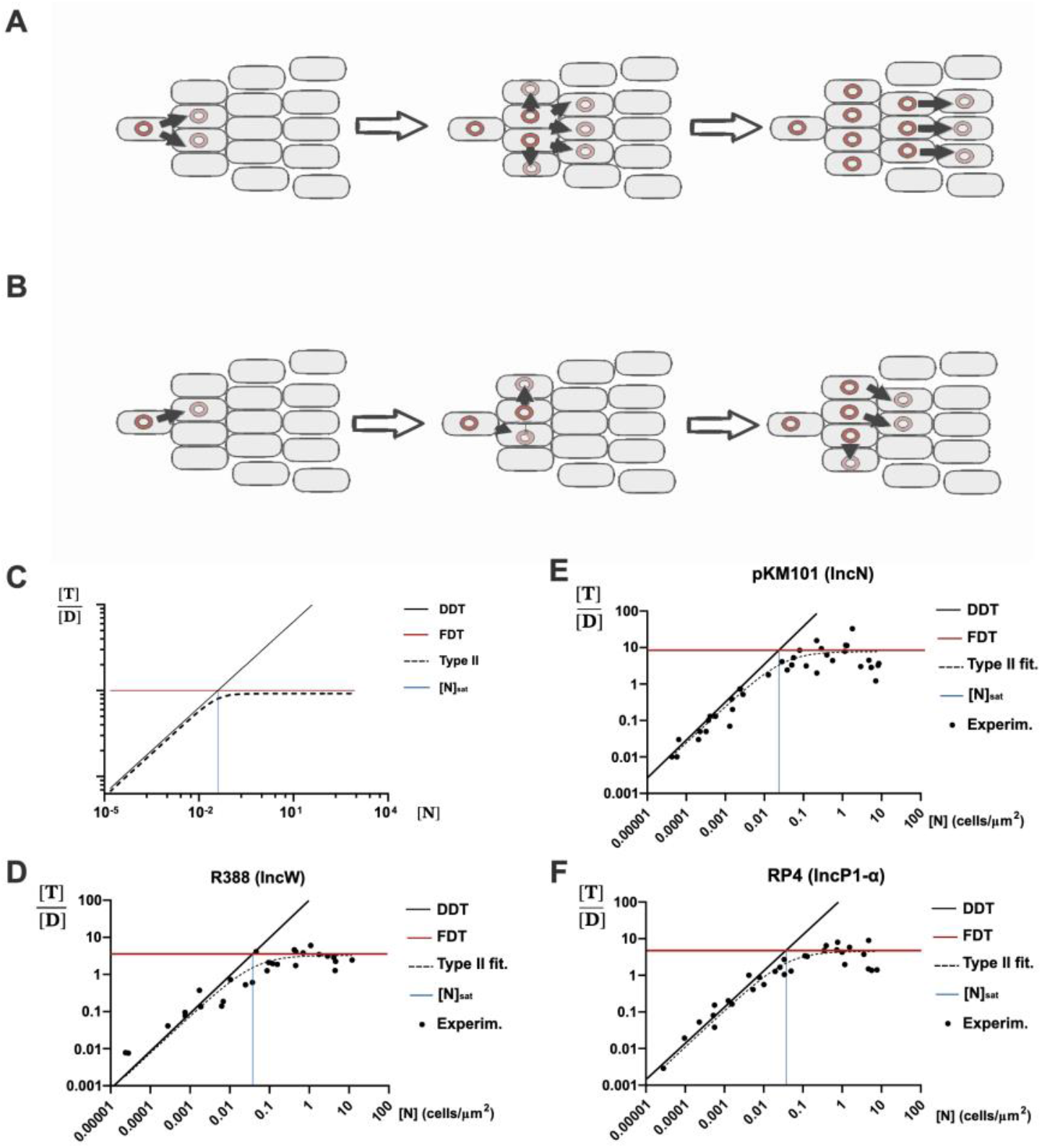
Conjugation follows a type II Functional response. **A**) UnderDensity-Dependent Transmission (DDT), plasmid conjugation increases linearly with increasingly available recipients; cell-to-cell contact is the only limiting factor, **B)** Under Frequency Dependent-Transmission (FDT), donors are limited by the time required to complete the transfer event, thus not all contacts result in successful matings. **C)** Ideal number of transconjugants obtained per donor (y axis, T/D) as a function of the total cell population (x axis, N). In a Holling’s Type II functional response there is a density-dependent region where transconjugant formation increases proportionally with N (leftwards from the blue line). When N becomes saturating the formation of transconjugants does not scale any further with cell density and becomes frequency-limited (rightwards from the blue line). **D, E, F)** Transconjugants per donor ([T]/[D], y axis) obtained at different cell densities ([N], x axis) in conjugations using E. coli BW27783, on solid LB-agar surfaces matings and 1 h. conjugation time. Each graph corresponds to plasmids R388 (D), pKM101 (E) and RP4 (F). The black line represents the ideal DDT dynamics, while the red line represents the FDT regime. The dotted curve corresponds to the fitting to Eq.3. Black dots correspond to the average of 3 replicates.

Here, by studying the dependence of plasmid transmission rates on recipient densities, we found that conjugation follows a Holling’s type II functional response. Plasmid transmission at a wide range of bacterial concentrations can be faithfully modeled from its K_on_ and τ values. These values were found to vary in different model broad host range plasmids, and even among members of the same plasmid taxonomic unit (PTU). Higher searching rates and lower handling times correlated with plasmid that displayed faster invasion dynamics. Overall, our results demonstrate that conjugation rates are intrinsically limited at a wide range of bacterial densities, but these limitations vary from plasmid to plasmid, suggesting that they may be genetically encoded and under active selection. Quantifying the transmission dynamics of different plasmids may shed light on the reasons for the different prevalence of PTUs in the microbiota, and their role in the spread of ARGs.

## Results

### Plasmid transmission becomes frequency-limited at moderate cellular densities

In a pure DDT regime, the number of transconjugants formed by each donor cell increases linearly with cell density (Figure 1C). In a pure FDT, however, the number of T/D cells obtained remains constant regardless of D and R concentrations (Begon et al., 2002). To analyse the transmission regime of conjugative plasmids we measured the T/D ratio exhibited by three model broad-host range plasmids with high conjugation frequencies: pKM101 (IncN/PTU-N1), R388 (IncW/PTU-W) and RP4 (IncP/PTU-P1). Conjugation experiments were performed between nalidixic (Nx) and rifampicin (Rif) resistant isogenic *E. coli* strains, as described in Materials and Methods. To minimize the confounding effects of vegetative growth, short mating times were employed (1h, unless otherwise stated), and the overall growth of D and R+T cells in these conditions was measured to be negligible (Supplementary Figure 1). To counteract the possible effects of structured populations, 1:100 and 1:10000 D:R ratios were used, so to ensure that each D cell is most likely surrounded by suitable recipients. In these conditions, results, shown in Figure 1.E-F, revealed that the number of transconjugants produced per donor cell scaled linearly with overall cell density, as expected for a DDT process, but reached a maximum at a density of approximately 0.05 cells/ μm^2^. This transition density, named [N]_sat_ for saturation density, roughly corresponds to 0.01 to 0.1 cells per square micron. From this density onwards, transmission followed an FDT dynamics, where conjugation rates were independent of the cellular concentration. Although the three plasmids switched from DDT to FDT regimes at approximately the same density, their transmission rates were different. Plasmid pKM101 showed a maximum rate of nearly 10 T cells per D per hour, significantly higher than that of plasmid RP4 (∼4 T/D*h) and plasmid R388 (∼3 T/D*h).

To evaluate whether this transition from DDT to FDT dynamics was characteristic of the rigid pili used to conjugate by the plasmids above, we repeated the experiment using plasmids pOX38 and R64 (IncFI/PTU-F1 and IncI1/PTU-I1, respectively). These plasmids use a flexible, long pilus to conjugate at high frequencies in liquid media (Bradley et al., 1980). As shown in Supplementary Figure 2, both pOX38 and R64 showed a similar behaviour to pKM101, RP4 and R388 plasmids, demonstrating that the shift from DDT to FDT occurs at similar densities, regardless of the MPF type harboured by the plasmid, when mating occurred on solid surfaces. To test whether this behaviour was also observed when conjugation was performed in liquid media, we analysed plasmid R64 transmission in LB broth. As shown in Supplementary Figure 3, T/D ratios saturated at cell densities >10^6^ cells/ml, indicating that this DDT / FDT transition also occurs in liquid media.

### Conjugation dynamics follows a type II functional response

Results from Figure 1. demonstrated that plasmid transmission did not follow pure DDT or FDT dynamics. To obtain an analytic model able to capture the entire dependence on cell density, we posed the following reaction scheme, where D, R and T correspond, respectively, to Donor, Recipient and Transconjugant cells.

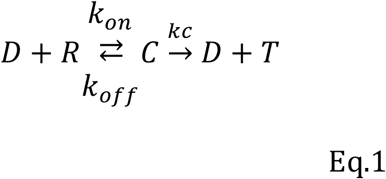

In Eq.1, D and R cells encounter each other following mass action kinetics, with a K_on_ rate. Formation of the conjugative pair, C, then ensues. C may resolve into the irreversible formation of a new transconjugant T at rate K_c_, or, alternatively, in an infructuous mating with a reverse rate K_off_. The handling time, hereafter τ, is the average time required for a donor to turn a receptor into a transconjugant, and can be expressed as 1/ K_c_. The model described in Eq.1 is suited for the transfer of mobilizable plasmids, where transconjugants are not able to become donors themselves. However, minor modifications allow the model to be adapted to self-transmissible conjugative plasmids (Supplementary Calculations). Eq.1 reaction scheme is conceptually identical to a Michaelis-Menten dynamics, and the same approximations can be used to obtain an analytic expression for transconjugant formation. As shown by Andrup and Andersen (Andrup & Andersen, 1999), following Haldane’s approximation, the number of transconjugants per donor (the conjugation frequency) obtained after time *t* follows:

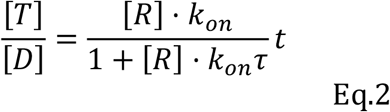

Eq.2 corresponds to Holling’s type II functional response, originally posed as the relationship between predation rate and prey density (Holling, 1959). In the context of plasmid conjugation, the density of susceptible recipient cells (R) divides the response into two distinct regimes (Figure 1C). At low R densities, the searching rate (k_on_) becomes limiting (density-dependent transmission). At high R densities, however, it is the handling time, τ, which limits plasmid propagation (frequency-dependent transmission). Holling’s type II functional response describes both regimes and the transition between them as cell density increases. Eq.2 is valid for plasmid mobilization, but as shown in Supplementary Calculations, plasmid conjugation can be similarly modelled. In this latter case, the autocatalytic nature of transmission (T cells become donors in conjugation, but not in mobilization) results into an exponential relationship of the form:

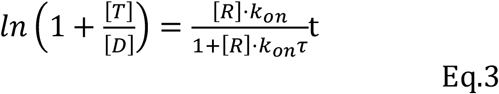

### Differences in conjugation parameters result in faster transmission dynamics

Data shown in Fig.1 was fitted to Eq.3 to extract the searching rate and handling times of plasmids RP4, pKM101 and R388. Results are shown in Table 1.

**Table 1.** Conjugation parameters obtained by fitting Eq.3 to data shown in Figure 1.

| Plasmid (PTU) | $K_{on}$ (Conf. interval at 95%) | $\tau$ (C.I. at 95%) |
| --- | --- | --- |
| R388 (PTU-W) | 90 $\mu\text{m}^2\cdot\text{h}^{-1}$ (45-130) | 0.33 h (0.25- 0.4) |
| RP4 (PTU-P1) | 160 $\mu\text{m}^2\cdot\text{h}^{-1}$ (115-200) | 0.25 h (0.2 - 0.3) |
| pKM101 (PTU-N1) | 215 $\mu\text{m}^2\cdot\text{h}^{-1}$ (180-250) | 0.13 h (0.1 - 0.2) |

Results indicated that pKM101 had higher searching rate K_on_ (*i*.*e*., donors carrying this plasmid found more available recipients during the same mating interval) and shorter handling times, τ (*i*.*e*., donors carrying this plasmid apparently need shorter times to successfully transfer the plasmid after contact), with respect to RP4 and R388.

To analyze the impact of differential searching rates and handling times on the overall kinetics of plasmid invasion, we monitored the progression of the plasmid in a susceptible *E. coli* population along time. Since plasmids pKM101 and R388 had previously shown the highest differences in handling times and searching rates, we performed invasion assays with these two model plasmids. As indicated in Materials and Methods, plasmid progression was measured as the number of transconjugant cells produced per donor cell (T/D). We performed these assays at two different D: R proportions, 1:100 and 1:10000, keeping the same total cell concentrations. Results, shown in Figure 2A, showed that both plasmids successfully invaded the population after 180 min., reaching T/D>10 in both cases. Plasmid pKM101 was significantly faster at both D:R ratios tested, demonstrating that higher searching rates and lower handling times result in faster plasmid propagation rates.

**Figure 2.**
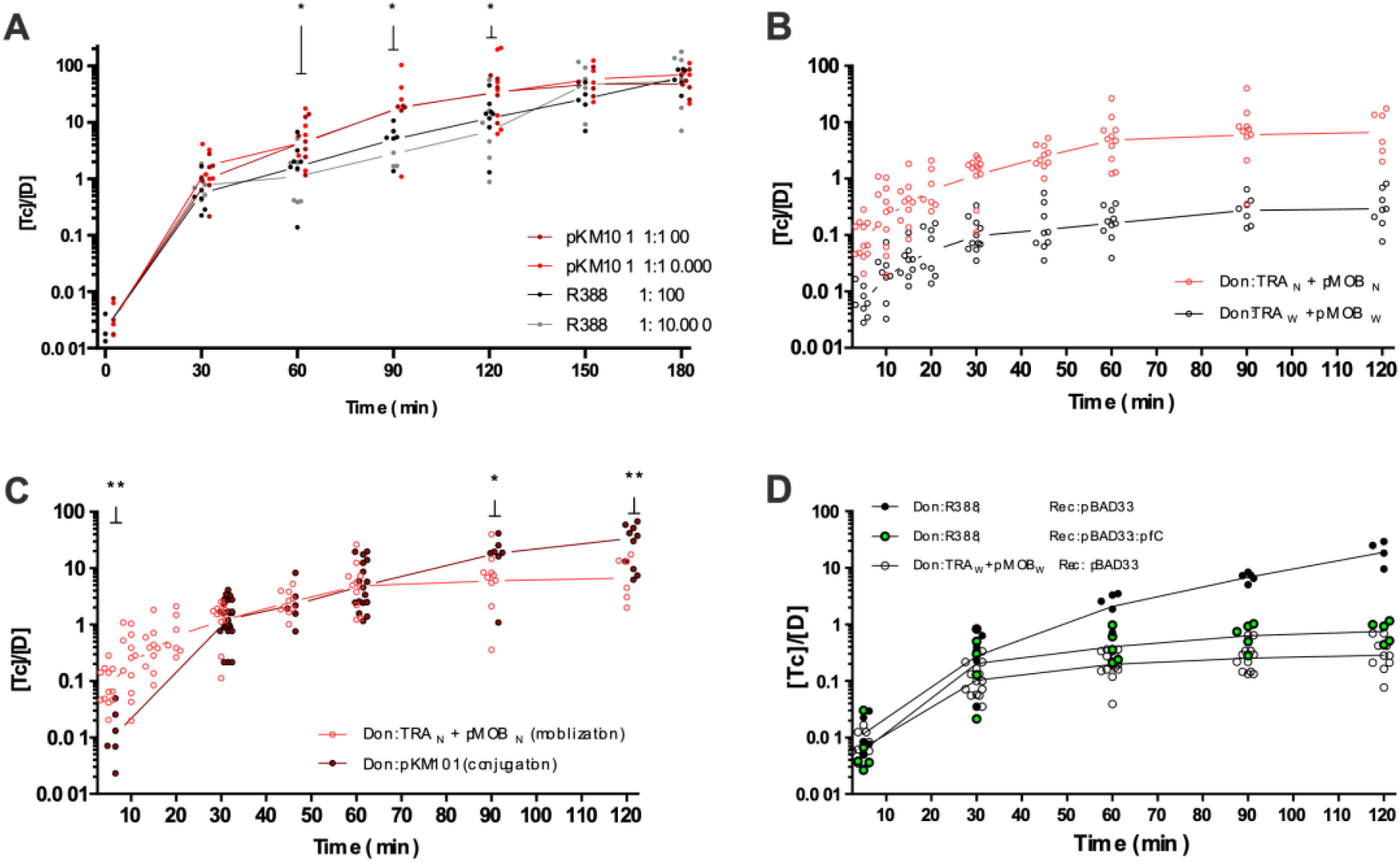
Plasmid invasion assays. Plasmid invasion of a susceptible population along time (x axis), expressed as transconjugants per donor (Y axis) **A)** Invasion dynamics of wild-type plasmids pKM101 (red dots) and R388 (black dots) obtained at two different donor: recipient ratios (1:100 and 1:10,000). Asterisks over t=60, t=90 and t=120 minutes indicate statistically significant differences between plasmids pKM101 and R388 with p<0.05 **B)** Mobilization dynamics obtained with TRA genes from PKM101 (TRA_N_, red) and TRA genes from R388 (Tra_W_, black) integrated in the chromosome of E. coli. The mobilizable plasmid contained the MOB region of the corresponding plasmid. Differences between plasmids were statistically significant with p<0.01 for all time points analysed **C)** Differences in the dynamics of conjugation (black) and mobilization (red) in plasmid pKM101. Mobilization assays were perform using a vector containing Mob_N_ harboured by a strain that contained the TRA_N_ genes inserted in the chromosome. Conjugation dynamics were obtained using the wild type pKM101 plasmid. Statistical differences were observed at t=90 (* p<0.05) and t=120 (** p<0.01) **D)** Comparison of plasmid R388 conjugation dynamics (black dots) vs R388-derived constructions. Black dots correspond to the wt plasmid transferred to a recipient strain that contains an empty pBAD expression vector. Green dots represent values obtained in conjugations where donors contained wild-type R388, and recipients harboured a pBAD::pifC construction, able to block secondary transmission. White dots correspond to the mobilization of a plasmid containing the MOB region of plasmid R388 from a strain harbouring a chromosomal integration of the Tra_W_ genes.

### Differences in the conjugation parameters are intrinsic to the conjugation machinery

Plasmid conjugation is a complex mechanism in which factors other than the transfer machinery often have an impact on the overall rate of plasmid propagation (Dionisio et al., 2005; Gama et al., 2017; Guynet et al., 2011). To test whether differences in searching rates (K_on_) and handling times (τ) were intrinsic to the conjugation machinery, or depended on the whole plasmid, we constructed a series of *E. coli* mutants harboring chromosomal constructions of the PTU-N1 or the PTU-W transfer systems. For this purpose, the entire TRA operon of each of the plasmids was introduced into the chromosome of *E. coli*, as indicated in Materials and Methods. These TRA_N_ and Tra_W_ constructions allowed the mobilization of plasmids encoding their cognate MOB regions (Mob_N_ and Mob_W_) to suitable recipients (Figure 2B). The overall T/D ratios obtained in mobilization assays were lower than those observed in conjugation, for both plasmids tested (Figures 2C and 2D). These differences could be due to the lack of secondary conjugation in mobilization assays. To test this end, we measured the conjugation dynamics of plasmid R388 when invading a recipient population expressing PifC. PifC is a protein that, when co-residing with the plasmid, blocks its transmission (Getino et al., 2017). This way, conjugation to a PifC-harbouring recipient should resemble plasmid mobilization, since the newly formed transconjugant is unable to transfer the plasmid in further conjugation rounds (Getino et al., 2017). As shown in figure 2D, the dynamics of plasmid R388 conjugation to PifC-containing recipients are similar to those of plasmid mobilization, suggesting that the differences in T/D ratios between conjugation and mobilization could be ascribed to lack of secondary conjugation in the latter.

A comparison between the mobilization rates of Tra_W_ and Tra_N_ (Figure 2B), demonstrated that the transfer machinery of pKM101 exhibited higher mobilization rates than R388. These results are consistent with the values obtained for wild type plasmids. Results in the searching rates and handling times observed in full plasmids are thus dependent on the differences between the transfer apparatus of both plasmids.

### Differences in conjugation rates are robust to environmental variables and overexpression of the transfer machinery

In some PTUs, environmental factors such as temperature, pH and nutrient availability have an important impact on plasmid transmission rates (Johnsen & Kroer, 2007). Thus, the differences observed between pKM101 and R388 could be caused by a differential sensitivity to environmental conditions. To test this end, we performed conjugations in different temperatures and nutrient conditions. Conjugation assays were performed as detailed in Materials and Methods, and results are shown in Figures 3A and 3B. As shown in Figure 3A, both plasmids exhibited an optimal transmission at 37ºC, yet pKM101 showed higher transfer rates at all temperatures tested. Plasmid conjugation rates were also relatively invariant when different media were used (Figure 3B): LB buffering against acidification (produced by carbon consumption) did not change the conjugation rates, as neither did increased nutrient availability by performing conjugations in terrific broth agar (TB agar). In all conditions tested, plasmid pKM101 consistently yielded higher conjugation efficiencies than R388.

**Figure 3.**
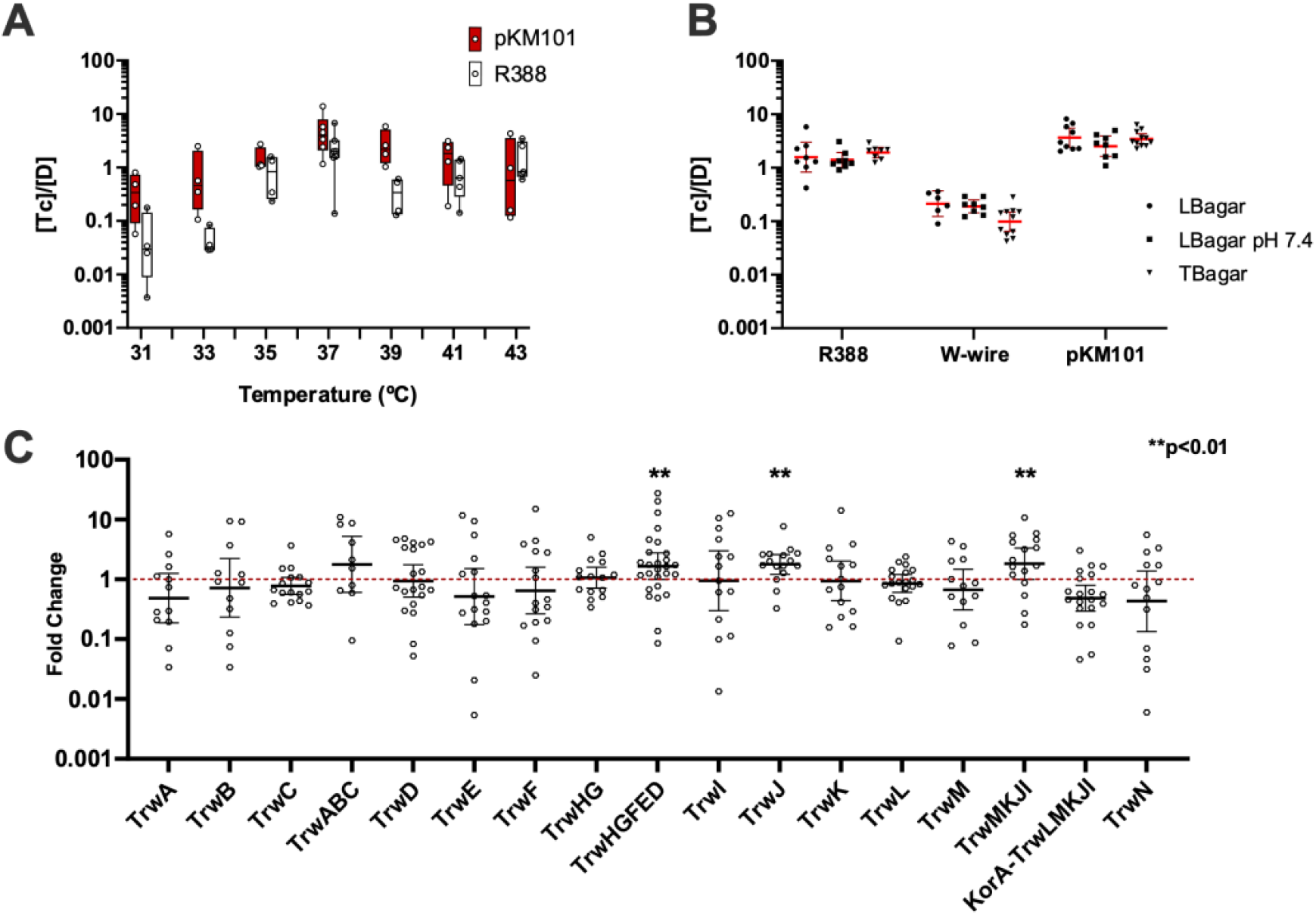
Effect of environmental conditions and protein expression on conjugation rates. **a)** Effect of temperature on the conjugation rate. Conjugation assays were performed as described in Materials and Methods. (1h, 37ºC, 1D:100R). Each point corresponds to an independent experiment **b)** Effect of media composition on the transfer rate for R388 conjugation and mobilization (W-wire) and pKM101 conjugation (LBagar: autoclaved LBagar, LBagar pH7.4: PBS-buffered at pH 7.4 and TBagar: Terrific Broth agar. **c)** Overexpression of TRA genes does not increase the transfer rate. On the x axis, genes overexpressed in R388-carrying donors using a pBAD33 vector. On the y axis, the fold change between the conjugation frequency (as [T]/[D]) obtained overproducing each of the constructions shown in the figure, and the conjugation frequency obtained with an empty pBAD33 vector. Statistical significancy was determined using Welch’s corrected ANOVA test for samples with unequal variances.

Another possible explanation for the differences observed between both plasmids was a differential expression of the conjugation apparatus. The expression of conjugation genes in both plasmids is regulated through two conserved negative feedback loops. The TRA operon is auto repressed by KorA, its first ORF, while MOB functions are controlled by second negative feedback under control of the accessory protein (TrwA_R388_/ TraK_pKM101_). Changes in these repressors or their cognate operators could result in different expression levels, thus we tested whether increasing the expression of conjugation genes altered plasmid transmissibility. Specifically, we analyzed whether increasing the expression of transfer genes increased the conjugation frequency of R388. For this purpose, MOB and TRA operons, as well as each individual conjugation gene, were cloned under the control of the pBAD promoter and introduced into donor cells. Overexpression of these genes was achieved as indicated in Materials and Methods, and the conjugation rates determined as previously described (Rodriguez-Grande & Fernandez-Lopez, 2020). Results, shown in Figure 3C revealed that the overexpression of transfer genes did not increase the conjugation frequency. Minor, yet statistically significative changes were achieved when overexpressing TrwJ, the whole TrwHGFED operon or the truncated TrwMKJI operon. TrwJ is the pilus adhesin, the protein responsible for contacting the recipient cell (Backert et al., 2008; Yeo et al., 2003). Thus, it is possible that overexpression of this protein results in better donor to recipient adhesion (Lacroix & Citovsky, 2011; Schmidt-Eisenlohr, Domke, & Baron, 1999; Schmidt-Eisenlohr, Domke, Angerer, et al., 1999, Frankel et al., 2023; Ishiwa & Komano, 2004; Low et al., 2023; Neil et al., 2020), increasing the apparent K_on_. The overexpression of the auto-repressed korA-TrwLMKJI operon led to decreased conjugation rates, probably due to overexpression of the korA repressor acting in trans on plasmid R388 transfer region (Fernandez-Lopez et al., 2014).

### Variation in the fundamental parameters within the PTU-W

Results demonstrated that pKM101 and R388 exhibited different conjugation parameters, which resulted in different invasion dynamics. These differences could be ascribed to their transfer machineries and were considerably orthogonal to environmental and transcriptional perturbations. We thus wondered whether the searching rate and the handling time were characteristic of a given plasmid PTU, or whether they were variable, with each plasmid showing its own, idiosyncratic values. To test this end, we compared plasmid R388 with three other PTU-W plasmids: pMBUI4, pIE321 and R7K. PTU-W plasmids have a highly conserved genomic backbone (Fernández-López et al., 2006), pIE321 being 97% identical at the DNA sequence level to R388, while R7K being 97.5% (a detailed comparison among them may be found in Revilla et al., 2008). Despite this high level of sequence conservation, these plasmids were isolated from different bacterial species. Plasmid pIE321 was isolated from *Salmonella enterica* serotype Dublin (Götz et al., 1996), R7K was found in *Providencia rettgeri* (Coetzee et al., 1972) and pMBUI4 directly from freshwater samples (Brown et al., 2013). Searching rates and handling times were obtained, as previously described, by analyzing the T/D obtained after 1h conjugation against a population of recipients with variable cell density (Figure 4A). While in pIE321 the handling time was slightly higher than that of R388, we observed a 10x increase in the case of plasmid R7K. To check whether these differences had an impact on the speed of propagation of R7K, we performed a conjugation kinetics assay, as described before (Figure 4B). As shown in the figure, the pace of spread in the recipient population was significantly slower in plasmid R7K, taking >150 minutes to reach a conjugation frequency of 1 T per donor cell. Plasmid R388 reached these levels in <30 min. Results thus demonstrated that within a given plasmid PTU, the conjugation parameters may exhibit significant variation, and this variation has an impact on the spread potential of the plasmid.

**Figure 4.**
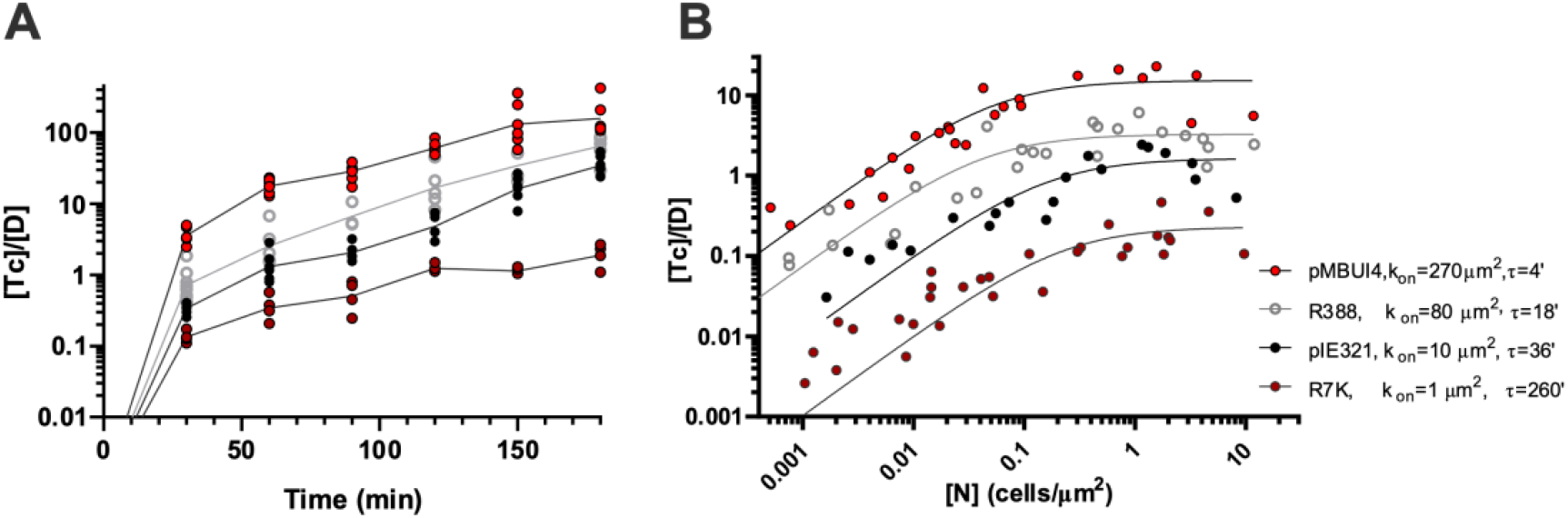
Plasmids from the same PTU display different conjugation dynamics. **(A)** Conjugation kinetics for different plasmids of the same PTU. The conjugation frequency, expressed as Tc / D is indicated on the y axis. The conjugation time, in minutes, is indicated on the x axis. **(B)** Conjugation frequencies (y axis) obtained at different cell densities (x axis) for different plasmids belonging to PTU-W. Estimated handling times (τ) and searching rates (k_on_) for each plasmid are indicated on the right side of the figure.

## Discussion

Understanding the factors that facilitate or curtail plasmid conjugation is essential to devise strategies against the propagation of antibiotic resistances. Plasmid propagation is a complex process, in which transmission, vegetative growth and plasmid-host co-adaptation play important roles (San Millan, A. & MacLean, R. C., 2017). Plasmid transmission is classically modelled as a mass-action kinetics process, but our results showed that this assumption breaks at moderate cellular densities of less than 0.1 cells per square micron. From this density onwards, conjugation resembles an FDT process, where the number of T cells produced per donor per unit time remains constant. By applying a model based on Holling’s type II functional response (Holling, 1959b), we were able to build a quantitative framework that faithfully recaptures the DDT/FDT hybrid regime. Our model depends on two parameters: the searching rate (k_on_, with units corresponding to area per time, μm^2^·h^-1^) and the handling time, τ (with units of time, h). The interplay between these two parameters (Eqs.2 and 3) determines the cell density at which density or frequency become limiting. By fitting conjugation frequencies obtained at different total cell densities, we were able to obtain estimates of both parameters for plasmids from the PTU-P1, PTU-N1 and PTU-W. Estimates of the searching rate, k_on_, were in all cases in the range of hundreds of square microns per hour, which roughly correspond to a searching radius of 6-10 µm^2^ per hour. These results are in accordance with estimates and recent direct observations of the effective distance for plasmid transmission, obtained for PTU-F_E_ and PTU-P1 plasmids (Harrington & Rogerson, 1990; Lagido et al., 2003; Babić et al., 2008; Seoane et al., 2011; Goldlust et al., 2023).

Interestingly, although both the searching rate and the handling time showed significant variation between plasmids, the cell density at which plasmids transitioned from one regime to the other was similar in all cases. Transition from a density-dependent to a frequency-dependent regime occurred at cell densities of approximately 0.02-0.05 cells per μm^2^ (that is, a 1 μm-long cell every 20-50 μm^2^). This figure roughly corresponds to a cellular density where each potential donor would be, at most, contacting 1 recipient. This is a surprisingly low cellular concentration, indicating that in denser environments such as biofilms (Fux et al., 2005; Hall-Stoodley et al., 2004; Madsen et al., 2012), soil (Raynaud & Nunan, 2014) or the gut microbiota (Contijoch et al., 2019) conjugation may be frequency-determined. This indicates that, in a significant proportion of microbial environments, plasmid conjugation is limited by the handling time, rather than the probability of encounters, as assumed by classical models of plasmid transmission (Levin et al., 1979; Simonsen et al., 1990; Smets & Lardon, 2009).

This frequency limitation explains a wide range of experimental results obtained for conjugation on solid surfaces and biofilms, which could not be explained by classical DDT models. In these experiments, a population of donors encountering one of recipients is typically able to generate only a “border” of transconjugants (Christensen et al., 1996; Fox et al., 2008; Reisner et al., 2012; Christensen et al., 1998; Hausner & Wuertz, 1999). Since handling times of conjugative plasmids are on the same order of magnitude as the doubling times of donors and recipients (Fernandez-Lopez et al., 2014), the layer of contact of recipients is pushed backwards at the same speed that the layer of transconjugants is created. When frequency becomes limiting, the factor constraining plasmid propagation is the time until the exhaustion of resources, compared to the handling time τ. The higher the number of effective transfers achievable by the plasmid before resources run out, the higher the conjugation rate. This explains why most plasmid propagation experiments on highly dense environments produced similar low invasion levels, except those continuously refreshing the growth medium (Turner, 2004) (Fox et al., 2008). Thus, to accurately measure the conjugative efficiency, experiments with short mating times (30-90’) and an excess of recipients (1:100 D:R ratio) are likely to yield more precise estimations. Results obtained in longer experiments, and with 1:1 D to R ratios would be confounded by vegetative growth and the lower probability of D-R encounters, respectively.

The handling time, τ, was found the most significant constraint for transfer. The factors that influence this parameter, however, are unclear. The fact that plasmids and their cognate transfer machineries exhibit slightly different handling times (Th=0.5h for pKM101 and Th=0.33h for Tra_N_) suggests that factors outside the conjugation region may play a role. The difference in size of the DNA being transferred is the most obvious candidate (pKM101 is 35 kb and the vector mobilized by Tra_N_ is approximately 5 kb). However classical experiments using HFR strains demonstrated that DNA transfer is fast, and the entire chromosome can be readily conjugated in approximately 100 minutes (Lederberg et al., 1952; Tatum & Lederberg, 1947). Other experiments also demonstrated that plasmid-encoded fluorescent proteins are expressed in transconjugants after 15 minutes of mating, indicating that the whole process of transfer takes shorter times (Gordon, 1992; Lawley et al., 2002). More recently, the timescale of the whole process of plasmid mobilization has been experimentally measured and seems to be in accordance with previous estimations (Couturier et al., 2023). Altogether, these results suggest that the handling time is not merely the time required to pump the DNA into the recipient. Instead, other factors conditioning the establishment and resolution of an effective conjugative pair may be at play. Plasmid copy number may be another possible contributor to τ. R388 and pKM101 plasmids are relatively low copied, compared to the vectors mobilized in assays using Tra_W_ and Tra_N_ mediated mobilizations. Moreover, the different IncW plasmids assayed in Figure 4 also present different copy numbers (Fernández-López et al., 2006), which could contribute to the differences observed in transfer rates. Further experiments systematically analyzing the impact of plasmid copy number on the transfer rate are required to clarify this end.

The searching rate K_on_, establishes how easily a successful D-R pair is established upon encounter. Physical factors involving cell to cell adhesion are likely to contribute this rate. It is known that conjugation is favored when other, unrelated conjugative plasmids express their pili in the same donor, or even the recipient cell (Gama et al. 2017). This suggests that increasing adhesion may facilitate transfer, a phenomenon that was also observed introducing synthetic adhesins in D and R cells (Robledo et al., 2022). Plasmids pKM101 and R388 showed a significant difference in their K_on_ (Table 1), a phenomenon which may be related to differences in the ability of the plasmid to facilitate cell to cell adhesion. Plasmid pKM101 contains *pep*, an adhesin-like protein that is absent in R388 and that has been shown to facilitate conjugation (González-Rivera et al., 2019). Also, the only protein that slightly increased the conjugation efficiency in R388 when overexpressed was the VirB5 adhesin TrwJ (Figure3). Similar results have been obtained in pKM101 (Yeo et al., 2003), pTi (Lacroix & Citovsky et al., 2011), and R64 (Komano, 1995), thus reinforcing the idea that increasing adhesion improves the conjugation rate. In fact, recent research shows that IncI and IncF plasmids selectively “pick” their Recipients via pilV or TraN-OMP surface interaction respectively (Allard et al., 2022; Frankel et al., 2023; Ishiwa & Komano, 2004; Low et al., 2023).

The fact that plasmids from the same PTU exhibited different searching rates and handling times was surprising. It indicates that, despite the overall sequence conservation, conjugation efficiency is not a conserved feature among the members of a PTU. In the case of W plasmids, sequence divergences concentrate in the MOB and replication regions, and are the highest between R388 and R7K plasmids (Revilla et al., 2008; Krol et al., 2013). Interestingly, R7K showed the lowest searching rate and the longest handling time, which resulted in slower invasion dynamics (Figure 4). R7K is the only PTU-W plasmid that was originally isolated from a host outside Enterobacteria, thus its poor performance may represent a maladaptation to *E*.*coli*, compared to the rest of PTU-W plasmids. Previous analyses have shown that plasmid-host co-evolution has a major impact on key plasmid properties such as host range and burden. (Bouma & Lenski, 1988; de Gelder et al., 2007, 2008; San Millan & MacLean, 2017). It is thus likely that plasmid transmission also depends on co-adaptations between the plasmid and the host. By systematically measuring the searching rates and handling times in plasmids at different stages of co-adaptation, we may be able to correlate transfer efficiencies with genes and mutations. Our model thus provides an analytic framework to unravel the mechanisms and constrains that regulate plasmid propagation in bacterial populations.

## Materials and Methods

Plasmids and strains employed in this work are detailed in Supplementary Tables 1 and 2.

### Cloning

Plasmid constructs used for gene dosage experiments were built using the Gibson assembly method (Gibson et al., 2009), using pBAD33 PCR products as vector and genes and operons from *tra* region of R388 as insert (to be inserted downstream the arabinose promoter of pBAD33 (Guzman et al., 1995)). All fragments were PCR amplified with Phusion DNA polymerase (Thermo Fischer), gel-purified and digested with DpnI restriction endonucleases (Promega) before assembly. *E. coli* DH5α strain was used for electroporation and selective culture purposes. Gibson assembly products were introduced in *E*.*coli* by electroporation. To build the mobilizable plasmid pSEVA321::mobW, the region containing *oriT* of RP4 included in vector pSEVA321 (coordinates 1278 to 1586 of GenBank Acc. No. JX560322.1) was replaced by the *oriT*-*trwABC* region of R388 (coordinates 11153-16299 of GenBank Acc. no. BR000038) using Gibson assembly.

### Tra_W_ and Tra_N_ constructions

The MDS42::Tra_W_ strain was constructed by cloning the *mpf* (also called *tra* operon) region of R388, comprising all genes (*trwD-korA*) required to build up the conjugative pili and inserting this whole region in a MDS42 *E*.*coli* strain (Pósfai et al., 2006) using Wanner-Datsenko protocol (Datsenko & Wanner, 2000). The lacZYA genes of E. coli strain MDS42 (coordinates 287438-292616 of GenBank Acc. No. NC_020518.1) were used as target for the chromosomal insertion. The trwD-korB region of R388 (coordinates 11153-16299 of GenBank Acc. No. BR000038) was PCR-amplified in three overlapping fragments. The kanamycin-resistance gene (KmR) surrounded by FLP recombinase recognition sites (FRT) was amplified from pKD4 (AY048743.1). Finally, 1kb fragments, homologous to the lacI and cynX chromosomal genes, were amplified from MDS42 genomic DNA. The six fragments were assembled (lacI-trwD-korB-KmR-cynX) and introduced by electroporation into MDS42 competent cells expressing the Lambda-Red site-specific recombinase (Datsenko & Wanner, 2000). KmR colonies were checked by PCR to confirm the Tra_W_ integration in the expected location, and the genome of a selected colony was completely sequenced by Illumina. The phenotype of the MDS42::Tra_W_ strains was checked by culturing in a chromogenic medium (agar Brilliance™) that differentiates lac- and lac+ phenotypes. The KmR cassette was subsequently deleted following the procedure described by Datsenko and Wanner (Datsenko & Wanner, 2000). Resistant derivatives of MDS42::Tra_W_ were obtained by plating saturated cultures of this strain in LB-agar plates supplemented with Rif10, Nx20 or Sm300.

MDS42::Tra_N_ strain was constructed by amplifying and Gibson-assembling the *mpf* region of pKM101 (genes *korB-traG*) in a pOSIP vector (St-Pierre et al., 2013), then electroporated in a DH5α *pir*^*+*^ strain before PCR comprobation, miniprep purification and introduction by electroporation in a *pir*^*-*^ MG1655 strain -following St-Pierre et al. protocol-where it recombined at attC position in the *E*.*coli* chromosome (St-Pierre et al., 2013).

### Conjugation and mobilization experiments

Unless otherwise stated, mating assays were performed using BW27783 (Khlebnikov et al., 2000, 2001) Nal^R^ (donor) and Rif^R^ (recipient) cells that had been grown to saturation overnight, at 37ºC, in 10 ml LB media supplemented with appropriate antibiotics. Antibiotic concentrations used were nalidixic acid 20 μg / ml, rifampicin 10 μg / ml, kanamycin 50 μg / ml, chloramphenicol 25 μg / ml, trimethoprim 20 μg / ml and ampicillin 100 μg / ml. Cells were washed in equivalent volumes of fresh LB without antibiotics and allowed to grow at 37ºC for 3h more. These cultures were collected, centrifuged at 4000g for 10’ and concentrated in 1 ml of fresh LB, then mixed at desired D:R ratios. Conjugation on solid surfaces were performed in Corning® Costar® 24 well plates to which 1 ml of LB-Agar had been added at least 3 days before, to solidify and dry. 15 µl of mating mix were placed on top of the LB-Agar and let to conjugate for 1 h. at 37ºC, except when indicated otherwise. Cells were then resuspended using 1 ml of sterile PBS, subject to serial dilution and plated on LB-agar with appropriate antibiotics.

Conjugation assays using serial dilutions to discern between FDT and DDT were performed as indicated above with the following modifications: For conjugation on solid surfaces, cell cultures of donor (10 ml) and recipient (100 ml) cells were grown to saturation on LB broth containing the appropriate antibiotics. Cells were washed, refreshed for 3h in LB without antibiotics, concentrated ∼100x in fresh LB and then mixed in 1:100 Donor to Recipient ratio before being serially diluted in LB (dilution ranging from 10^2^X to 10^−4^X). From each dilution, 100 μl of cells were deposited on top of a 10cm diameter LB-agar plate (also prepared at least 3 days before to ensure proper LB agar drying) and incubated for 1h at 37ºC. Cells were resuspended in 1000 μl of PBS, serially diluted and appropriate dilutions were plated on LB-agar plates with antibiotics to count the number of D, R and T cells present.

Every D, R and T count corresponded to three technical replicates. Cell densities (in cells/µm^2^) were calculated dividing the average of these 3 counts per total surface: 2 cm^2^ in 24-well plates, 58 cm^2^ in Petri plates.

The same protocol was applied in mating experiments at different times, with only two changes: a 1:10.000 ratio was also used (by putting 1/100 less donor volume in the same amount of 10x concentrated recipient cells, as usual) and PBS was used to abort conjugation at different times: 30, 60, 90, 120, 150 and 180 minutes. Several methods for measuring this efficiency have been used (Dimitriu et al., 2019; Dionisio et al., 2002; Gama et al., 2017; Inoue et al., 2005; Kosterlitz et al., 2022) and discussed elsewhere (del Campo et al., 2012; Simonsen, 2009; Simonsen et al., 1990; Wan et al., 2011; Zhong et al., 2012), but we will use T/D as a measurement of conjugation (or mobilization) efficiency on surfaces because at very short times (<3h) it is free from factors like differences in growth or population structure, still depicting very intuitively the differences in transmission efficiency, and its underlying causes.

Experiments measuring gene dosage effect on conjugation were conducted as the rest of conjugation experiments explained above, except for the donor and recipients were grown overnight on LB with antibiotics and glucose 0.5% (for pBAD33 arabinose promoter repression) and washed and allowed to grow for 3h more in LB without antibiotics with glucose 0.5% or arabinose 50μM (for arabinose promoter induction). Each experiment was conducted simultaneously with a control donor containing the conjugative plasmid and pBAD33 with no cargo. No transfer of pBAD33-derived plasmids was observed. Conjugation efficiencies were determined as T/D and ratios mean conjugation efficiency (three replicates) for the plasmid + pBAD33::trwX donors divided by mean conjugation efficiency (three replicates) for the plasmid + pBAD33 control were represented in graphs. In this case, if no change in conjugation efficiency is observed, the expected mean ratio clone:control should be 1, while if efficiency is improved by supplementing a gene or operon, the ratio should be greater than 1. In this case, DH5α strain was used as donor to avoid recombination between vector cargo and R388wt.

Conjugation experiments in buffered LB were performed at a pH=7.4 using Na_2_HPO_4_ 10mM and NaH_2_PO_4_ 1.8mM and pH adjusted with HCl 35% and NaOH 1M. Conjugation experiments using filtered LB agar were performed as described, but using LB medium filtered with a 0.22 μm filter membrane instead of using high-pressure sterilized LB. On the same manner, TB agar experiments were performed as usual, but using autoclaved Terrific Broth instead of LB, with the same medium to agar ratio.

### Statistical tests

In order to check if conjugation efficiency measurements (as T/D cell counts) did follow a normal distribution we performed a Shapiro-Wilk and an Anderson-Darling test on two independent arrays of 16 measurements of R388 conjugation efficiency each. Both groups of T/D counts showed no normality, but when the analysis was performed on their log-transformed values they showed normal distribution (Shapiro-Wilk, W=0.97 both groups, Anderson-Darling test A*^2^=0.26 and 0.21 respectively), thus indicating log-normal distribution, as suggested before (Gama et al., 2017; Lucas et al., 2010).

Paired comparisons of log(T/D) values were performed using two-sided t Student’s test, while multiple comparisons were performed using ANOVA. Unless otherwise specified, a significance level of α= 0.05 was taken as good enough to reject null hypothesis.

Nonlinear fitting of conjugation efficiencies at different cell concentrations was performed with nonlinear curve fitting package of GraphPad Prism v8.0.2 using custom equations (namely, Eqs 2 and 3).

## Acknowledgements

This work was supported by the Spanish Ministry of Science and Innovation (Project PCI2021-122067-2A funded by MCIN/AEI/10.13039/501100011033 and the European Union NexGenerationEU/PRTR and by Project PID2019-110216GB-I00 funded by MCIN/ AEI /10.13039/501100011033 to RFL)

---

**Supplementary Figure 1.**
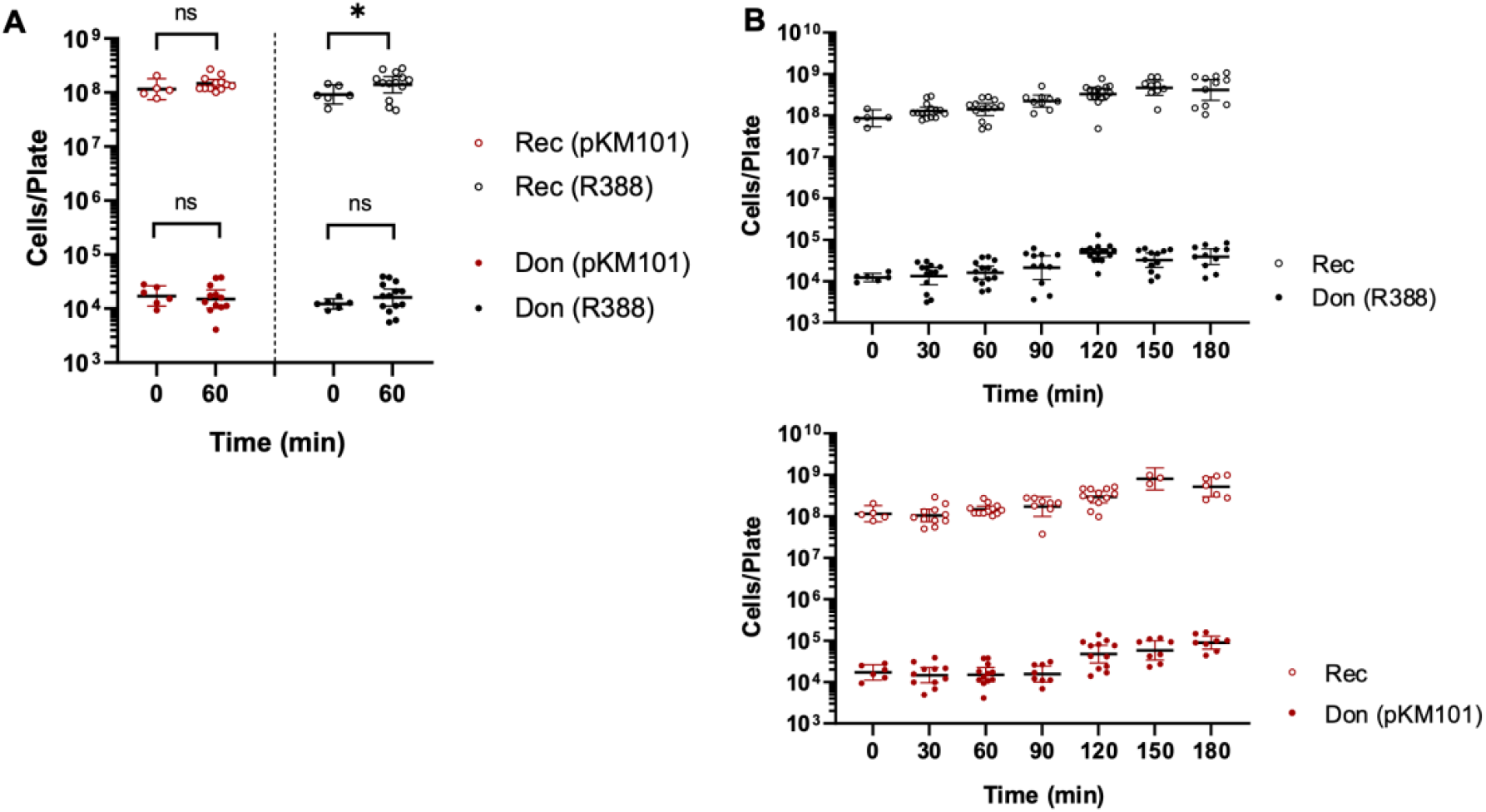
A) Growth of Donor (solid circles) and Recipients + Transconjugants (hollow circles) during the first hour of mating in conjugation experiments of R388 (black, right) and pKM101 (red, left) carrying cells at 1:10,000 D:R ratios as detailed in Figure 2A in the main text. ANOVA testing comparing the four populations at 0 and 60 minutes after mating began gave p>0.05 in all but R388 Recipients’ growth. B) Growth curves for donor and recipients in matings using plasmid R388 (black circles, upper graph) and pKM101 (red circles, lower graph) fitting showed that doubling times were >60 minutes for all strains analyzed.

**Supplementary Figure 1.**
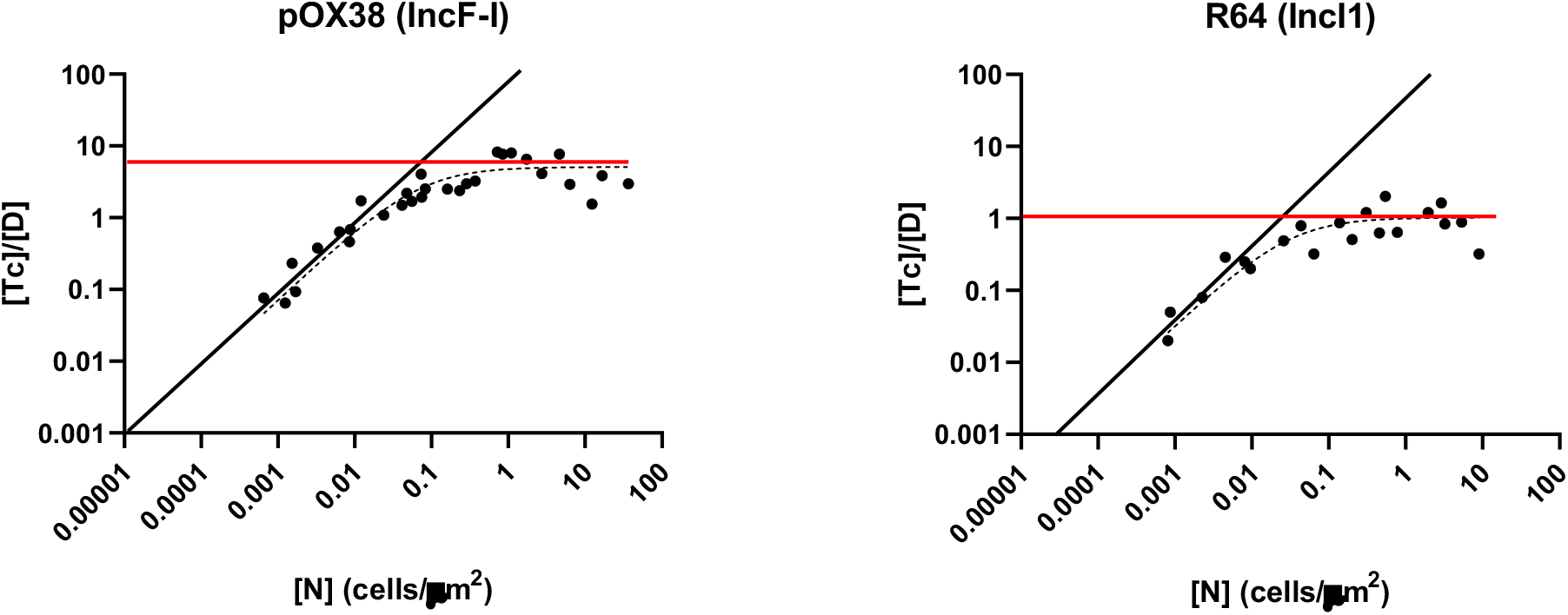
DDT / FDT transition in conjugations with plasmid pOX38 and R64 on solid media. Transconjugants per donor ([T]/[D], y axis) measured at different Recipient densities ([R]*≈*[N], x axis) in E. coli BW27783 mating on solid LB-agar surfaces, 1 Donor to 100 Recipients and 1 h. conjugation time for plasmids pOX38 (IncFI) and R64 (IncI1). As in Figure 1 in the main section, the black line represents the ideal DDT regime, where different *k*_on_ produce different y-intercepts. The red line represents the FDT regime, where the conjugation efficiency is just 1/ τ and the dotted curve corresponds to the fitting to Eq.3. Every black dot corresponds to the average of 3 technical replicates.

**Supplementary Figure 3.**
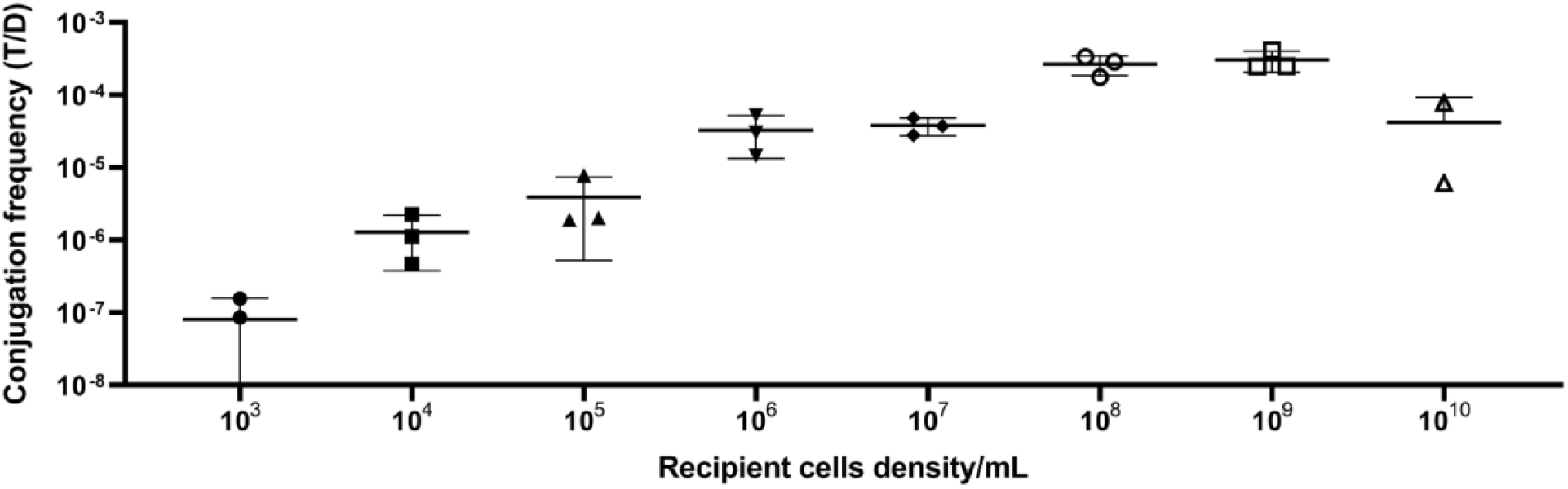
DDT / FDT transition in conjugations with plasmid R64 in liquid media. Transconjugants per donor ([T]/[D], y axis) measured at different Recipient densities ([R]*≈*[N], x axis) in E. coli BW27783 mating in liquid LB, 1 Donor to 100 Recipients and 1 h. conjugation. Every dot corresponds to the average of 3 technical replicates.

**Supplementary Table 1.** Strains used in this work.

| Strain | Genotype | Reference |
| --- | --- | --- |
| **DH5α** | F<sup>-</sup> endA1 hsdR17 (rk<sup>-</sup>, mk<sup>+</sup>) supE44 thi-1 λ<sup>-</sup> recA1 gyrA96 relA1 deoRΔ(lacZYA-argF)-U169 φ80d lacZΔM15 | (Grant et al., 1990) |
| **BW27783** | lacI<sup>q</sup> rrnB<sub>T14</sub> lacZ<sub>WJ16</sub> hsdR514 Δ(araBAD<sub>AH33</sub>) Δ(rhaBAD<sub>LD78</sub>) Δ(araFGH) φ(ΔaraEp P<sub>CP8</sub>-araE) | (Khlebnikov et al., 2000, 2001) |
| **MG1655** | Dam<sup>+</sup>, F<sup>-</sup>, lambda<sup>-</sup>, rph-1 | (Blattner et al., 1997) |
| **MDS42** | MG1655Δ(fhuACDB) endA<sup>-</sup>, recA<sup>-</sup>, Δ(699 genes, ISs and prophages) | (Pósfai et al., 2006) |
| **MDS42 Tra<sub>N</sub>** | MG1655::mpf_R46 | This work |
| **MDS42 Tra<sub>W</sub>** | MDS42::mpf_R388 | This work |
| **DH5α pir<sup>+</sup>** | endA1 hsdR17 glnV44(=supE44) thi-1 recA1 gyrA96 relA1 φ80dlacΔ(lacZ)M15 Δ(lacZYA-argF)U169 zdg-232::Tn10 uidA::pir<sup>+</sup> | (Platt et al., 2000) |

**Supplementary table 2.**
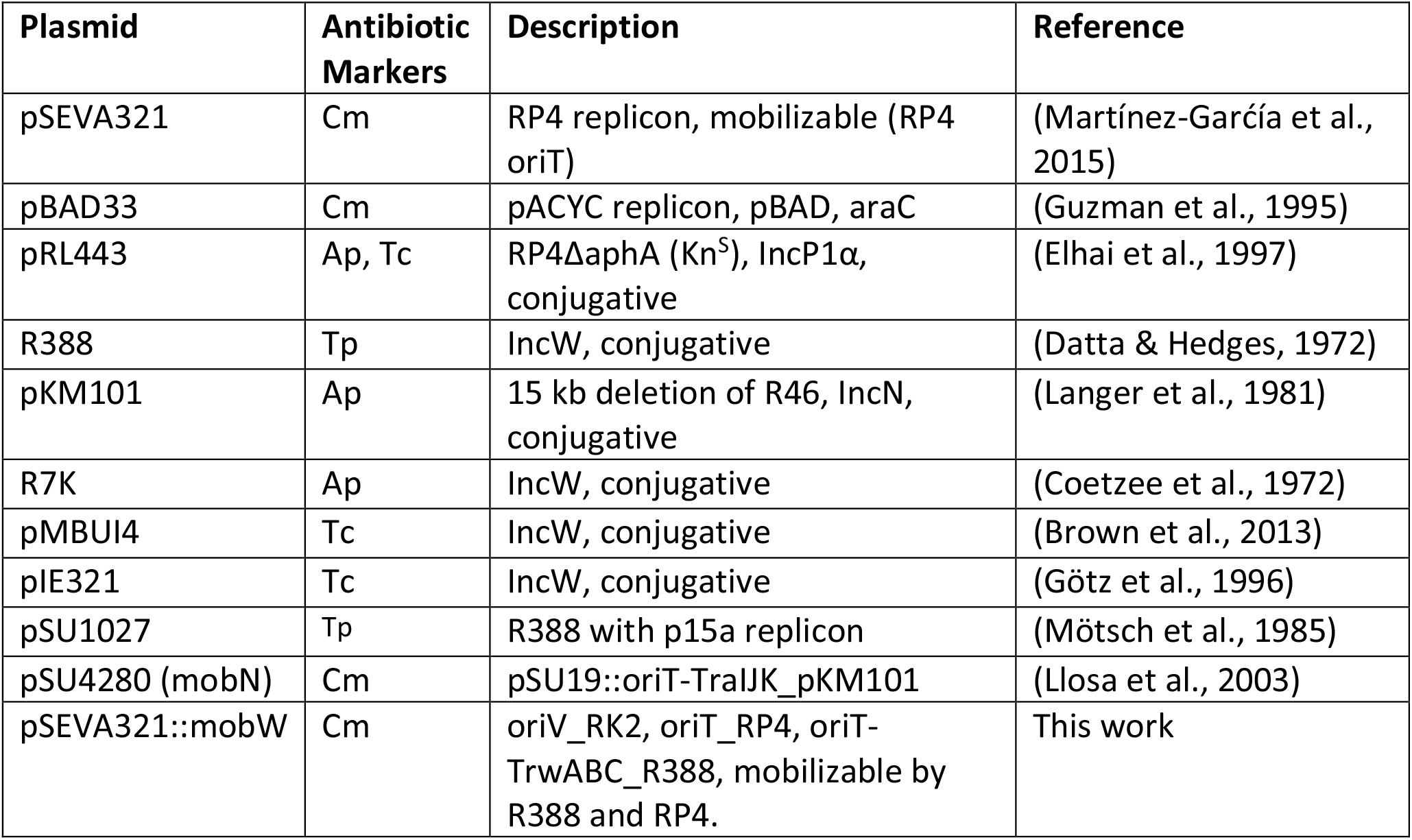
Plasmids used in this work.

## SUPPLEMENTARY CALCULATIONS

### 1. Deriving the functional response for plasmid mobilization

To obtain the dynamics of mobilization we pose a model with three different types of cells:

*D* = donors

*R*= recipients

*T*= transconjugants

The model assumes that *D* cells encounter *R* in a density-dependent fashion, producing a complex *C*. The conjugative complex may progress to the formation of a transconjugant *T*, or may result in a non-productive mating (by premature termination of the conjugative pair), such that:

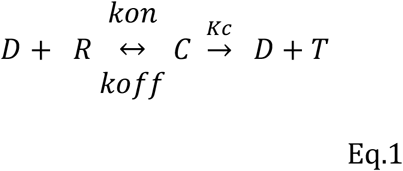

We also will perform our experiments in the regime where *R* >> *D*, thus the number of recipients may be considered a constant. Under these assumptions, the differential equations governing the progression of the population follow:

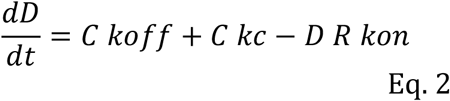

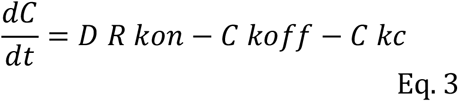

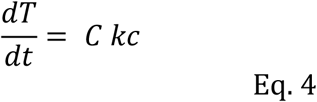

The system is thus formally equivalent to a Michaelis-Mentem reaction scheme, following the Haldane approximation, we can pose the following:

1) Pseudo-steady state: Given enough time *t*, formation of *C* complexes will be in steady-state, thus:

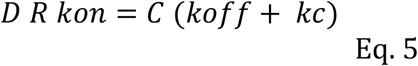

2) Preservation of the total number of *D*: the contribution of vegetative growth during the experiment is considered negligible

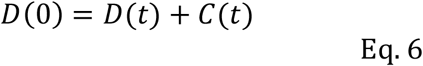

This allows us to pose an equation which is formally equivalent to that of the MM process under the Haldane approximation:

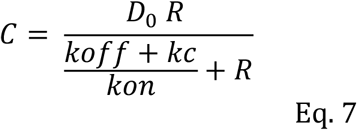

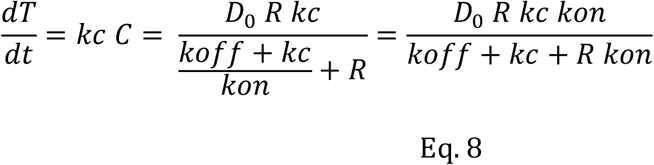

Now, we can introduce the average lifetime of C, the time required for the C conjugative complex to resolve into the formation of transconjugants or an abortive reversion to D + R. Such average lifetime (τ)is just 1 / (*koff* +*kc*). Introducing this factor into Eq.8 yields

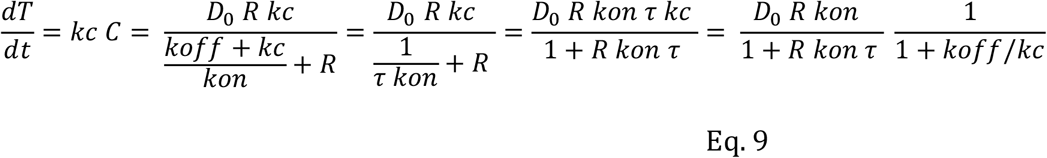

This Eq can be now directly integrated yielding the following expression for the formation of transconjugants per donor :

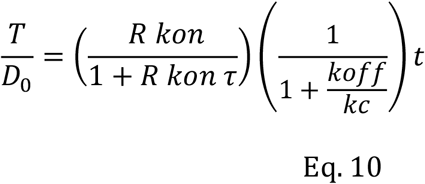

Now, in conditions where the formation of a transconjugant is much more likely than the abortion of the conjugative pair, such that kc>> koff Eq 10 simplifies to an expression which is formally equivalent to Holling’s type II functional response:

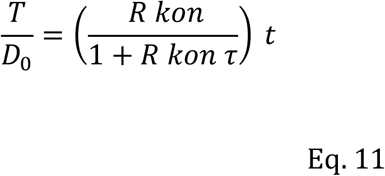

In this Eq the transconjugants generated per donor cell depend on two constants: a *kon* parameter that indicates the rate at which donor cells find and attach to recipients, and an effective conjugation time, *τ*, indicating the time required for a donor-recipient complex to resolve into a new transconjugant. In those cases where the abortion rate of the conjugative plasmid is substantial, the rate of transconjugant formation will be affected by a term which is just the probability of the C complex resulting into the successful formation of a new transconjugant:

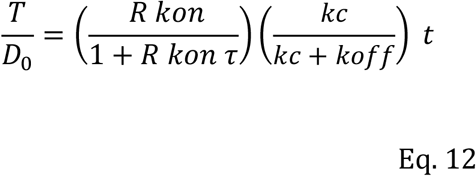

### 2. Deriving the functional response for plasmid conjugation

The dynamics of plasmid conjugation are significantly more complex than that of mobilization. The newly formed transconjugants can act as donors, a process which is often assumed to involve a lag period. Additionally, transconjugants often exhibit higher conjugation rates than the parental donor population, due to the transitory de-repression of the conjugation machinery after plasmid transmission. An autocatalytic process of this nature, involving lags of variable time and non-linear dynamics, frequently exhibits too complicated dynamics to be faithfully parametrized. We can, however, modify the experimental conditions to simplify the dynamics and obtain the searching rate and the effective conjugation time.

The simplest approximation is to assume that, in conjugative plasmids, the rate of formation of new transconjugants is proportional to the number of donors D_0_ and already formed transconjugants (T_t_). In these conditions:

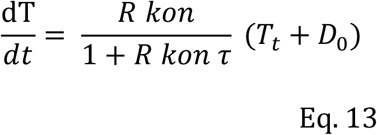

This expression can be directly integrated, yielding:

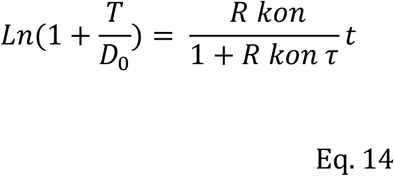

For plasmids with low conjugation rates the overall contribution of the newly formed transconjugants is negligible, compared to that of the initial D population. This translates in the convergence of Eq 14 and Eq 11 at low conjugation frequencies. As a rule of thumb, in a plasmid with a conjugation frequency of 0.1 T/D the contribution of secondary conjugation is approximately 0.01 T/D. This means that for most experimental conditions and model plasmids, Eq10 can be directly applied without the introduction of a substantial estimation error. This, however, does not hold for plasmids in which the conjugation frequency observed is higher than 1 T/D (for 1 T/D the error would be 0.3 T/D). This may be the case of highly conjugative plasmids or, more frequently, mating experiments performed for very long times, under a continuous replenishment of nutrients.

### 3. Vegetative growth contribution is negligible

In Eq.12 we assumed vegetative growth to yield a negligible contribution to the frequency of conjugation (Fc). Here we will show that this is always the case, as long as the growth rate of T and D is similar.

The FC is defined as the ratio of T to D, such that

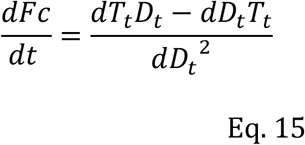

Where

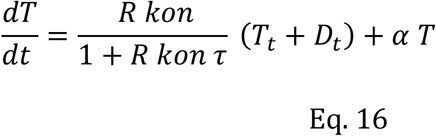

And the growth of *D* follows:

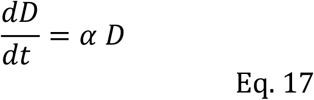

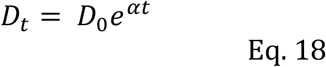

In these conditions

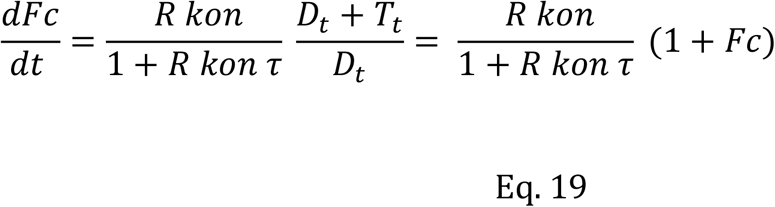

Which can be directly integrated

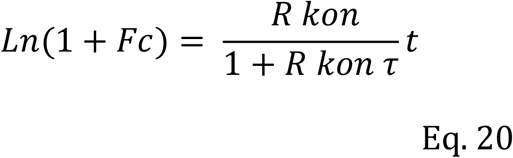

